# Mutations in components of the TREX-2 complex result in misexpression of the Kelch-domain F-Box protein KFB39 promoter in *Arabidopsis thaliana*

**DOI:** 10.1101/2025.10.22.683924

**Authors:** Michaela C. Matthes, Smita Kurup, John A Pickett, Johnathan A. Napier

**Affiliations:** Plant Sciences, Rothamsted Research, Harpenden, Herts AL5 2JQ, UK

## Abstract

*Arabidopsis thaliana* gene At2g44130 encodes a Kelch-like domain F-box protein designated KFB39 and was previously shown to be specifically expressed on exposure to the oxylipin *cis*-jasmone. In order to better understand the regulation of At2g44130, a forward genetic screen was carried out to identify mutants in which a promoter-GUS fusion was expressed in the absence of the inducer, *cis*-jasmone. Two mutants were recovered, showing mis-expression of the promoter-GUS fusion and surprisingly both were found to be in components (SAC3B, THP1) of the TREX-2 nuclear pore complex. Genetic analysis of *sac3* mutants in Arabidopsis revealed additive impairments to growth and development as well as reduced capacity for nuclear export. Promoter-GUS fusions of the Arabidopsis SAC3 and THP1 genes revealed a discrete expression pattern that was non-overlapping with KFB39. A link between the expression of KFB39 and the TREX-2 complex is not obvious, but we note that previously, unrelated forward genetic screens using promoter-reporter fusions have also recovered *sac3b* and *thp1* mutants. We consider some possible explanations for these shared occurrences.

## Introduction

Previously, we identified a suite of Arabidopsis genes upregulated by the hyper-volatile oxylipin *cis*-jasmone (Matthes et al., 2010). *cis*-jasmone (hereafter abbreviated to cJ) is often considered to be a catabolite of jasmonic acid (Dabrowska & Boland 2007) but it is now clear that it plays many critical functions in plant biology beyond just being a disposal route for the attenuation of the jasmonate signal, most notably in plant-insect interactions (Bruce et al., 2008). In addition, the reactive electrophile nature of cJ means that it generates a distinctive transcriptome to that induced by jasmonic acid or methyl-jasmonate (Koster et al., 2012). In that respect, it is likely that cJ is partially perceived as a xenobiotic insult, emphasised by the COI1-independent expression of many cis-jasmone-induced genes which at the same time were shown to require SCL and TGA transcription factors (Mueller et al., 2008; Matthes et al., 2010). Of all the Arabidopsis genes which were induced by cJ, one was of particular interest – At2g44130, a Kelch-like domain F-box (KFB39) protein, which was strongly induced by cJ but independent of COI1 and JAR1 as well as TGA2,5,6 and SCL14 (Matthes et al., 2010), confirming it as distinct from both the jasmonate and reactive electrophile signalling pathways (Farmer & Mueller, 2013). Promoter-reporter (GUS) fusions of the upstream sequence of At2g44130 revealed a discreet tissue-specific expression pattern for this gene in the presence of cJ, predominantly in the midrib of the leaf but otherwise transcriptionally silent in the absence of induction from this chemical signal (Matthes et al., 2010; Curtis et al., 2013).

To better understand the factors that determined the expression of this gene, as well as to shed additional light on the regulation of gene expression by xenobiotic factors, we carried out a forward genetic screen to isolate mutants in which the GUS reporter was expressed from the At2g44130 promoter in the absence of the inducer cJ. By this means, we hoped to identify upstream genes involved in the expression of the KFB39 F-Box protein. Here we report the unexpected identification of two different components of the TREX-2 nuclear pore complex and the possible role for this complex in the regulation of the KFB39 gene. We also consider how our observations fit with other studies in which alleles of the same components of TREX-2 have been isolated from unrelated forward genetic screens in Arabidopsis.

## Materials and Methods

### Plant material

We used the Col-0 ecotype *of Arabidopsis thaliana* for the studies reported here. T-DNA KO lines for SAC3B (Salk_111245 and Salk_065672), SAC3A (SAIL_584_A02), SAC3C (Salk_083805) and THP1 (SAIL_82_A02) were obtained from NASC. Besides the two lines harbouring the knock-out alleles for SAC3B all other lines were previously described by Lu et al., 2010. Plants were grown in a Weiss Gallenkamp cabinet under long day conditions (16h light 23°C/ 8h dark 18°C) and 250 μmol m^-2^ sec^-1^ white light provided by a fluorescent lamp.

### Poly(A) RNA *in situ* hybridization

Poly(A) RNA *in situ* hybridization was performed essentially as described by Gong et al., 2005. In brief, samples were taken from equivalent portions of young leaves at similar developmental stages from the wild type and mutants and were fixed in glass vials by adding 10 mL of fixation cocktail containing a mixture of 50% fixation buffer (120 mM NaCl, 7 mM Na_2_HPO4, 3 mM NaH_2_PO4, 2.7 mM KCl, 0.1% Tween 20, 80 mM EGTA, 5% formaldehyde, and 10% DMSO) and 50% heptane. The samples were gently shaken for 30 min at room temperature. After dehydration twice for 5 min in absolute methanol and three times for 5 min in absolute ethanol, the samples were incubated for 30 min in 1:1 ethanol:xylene and then washed twice for 5 min with absolute ethanol, twice for 5 min with absolute methanol, and once for 5 min with 1:1 methanol:fixation buffer without 5% formaldehyde. The samples were postfixed in the fixation buffer for 30 min at room temperature. After fixation, the samples were rinsed twice with fixation buffer without 5% formaldehyde and once with 1 mL of PerfectHyb™ hybridization buffer (Sigma-Aldrich; H-7033). Each glass vial was then added with 1 mL of hybridization buffer and prehybridized in an incubator for 1 h at 50°C. After prehybridization, 5 pmol of 45-mer oligo(dT) labelled with one molecule of fluoresceine at the 5⍰-end (synthesized by Sigma-Aldrich) was added into each glass vial and hybridized at 50°C in darkness for more than 8 h. After hybridization, the samples were washed once for 60 min in 2× SSC (1× SSC is 0.15 M NaCl and 0.015 M sodium citrate), 0.1% SDS at 50°C and once for 20 min in 0.2× SSC, 0.1% SDS at 50°C in darkness. The samples were observed immediately using a ZEISS 780 Confocal Laser Scanning Microscope (ZEISS, Germany) with a 488-nm excitation laser and a 522/DF35 emission filter. All samples were observed at the same conditions and the experiment was repeated three times with similar results.

### Ethylene assay

Ethylene response of WT, mutant and KO lines was evaluated by measuring hypocotyl length of seedlings after 3 days of growth in the dark on half-strength MS medium with or without 50 μM of the ethylene precursor ACC. Images of seedlings were taken with a Leica M205 FA stereomicroscope and hypocotyl length measured with ImageJ.

### GUS assays

To construct the Sac3B:GUS reporter fusion 0.9 kb of the SAC3B (At3g06290) promoter were amplified with primers Sac3BHindIII (5’-AAGCTTCGCCATTATCGTTATCAGCAACC-3’) and Sac3BNcoI (5’-CCATGGTCTGAGAAAATTCCTTGACTTG-3’) and the fragment inserted into pJD330 from which the CaMV 35S promoter had been excised by HindIII and NcoI. The promoter:GUS reporter fusion was subsequently cloned into BIN19 using the restriction enzymes HindIII and EcoRI. A 1.27 kb fragment of the Sac3A promoter (At2g39340) was amplified with primers Sac3AKpnI (5’-GGTACCATCCTACCCTACTTGCTCATAAAC-3’) and Sac3ANcoI (5’-CCATGGTTTATCATCAACGTCCTGAAAA-3’). The fragment was subsequently inserted *via* its KpnI and NcoI restriction sites into a pBS plasmid which contained the GUS-nos terminator sequence from pJD330 cloned by SalI and EcoRI (pBS GUSpJD). The Sac3A promoter:GUS fusion was excised using KpnI and PstI, the fragment blunted and cloned into the SmaI site of BIN19. 0.92 kb of the Sac3C (At3g54380) promoter were amplified with primers Sac3CSalI (5’-GTCGACCAGAGAATGCTTGGTAGATAG-3’) and Sac3CXhoI/KpnI (5’-GGTACCCTCGAGGGCTTAGAGGGAACGAGATCTTCTT-3’) and subcloned into pCR®-BluntII-TOPO (Thermo Fisher Scientific). Subsequently, the promoter fragment was cloned into BIN19 using the SalI and KpnI sites. The GUS-nos terminator sequence was excised from pBS GUSpJD by XhoI and EcoRI digestion and inserted into the BIN19 vector containing the Sac3C promoter which had been cut with XhoI and EcoRI. Constructs were transferred to Arabidopsis by floral dip. Per construct several independent lines were used for GUS staining. Histochemical characterization was carried out by incubating tissue in a solution containing 5-bromo-4-chloro-3-indolyl glucuronide (Melford, UK), 0.1M KH_2_PO4 buffer [pH 7], 0.1% Triton X-100, 5 mM potassium ferrocyanide and 5 mM potassium ferricyanide. Vacuum was applied to the tissues briefly before incubation at 37°C overnight in the dark. Tissue was subsequently cleared in 70% ethanol. Images were taken either with Leica M205 FA stereomicroscope or a ZEISS Axioimager.

## Results

### Isolation and identification of mutants de-regulated in the cis-jasmone induced expression of the Kelch-like domain F-box KFB39

An Arabidopsis Col-0 line expressing the GUS reporter under the control of the At2g44130 promoter (pF-Box:GUS) was subjected to mutagenesis with EMS following standard protocols – as described reported by us, this promoter-reporter fusion is inactive in the absence of the oxylipin inducer cJ (Curtis et al., 2013). Pools of 25 plants per pool of the mutated population were screened for constitutive expression of GUS in the absence of cJ, comprising a total of 2500 plants. We found that two lines (designated 48_67 and 89_75) mis-expressed the reporter and displayed a distinctive pattern of GUS accumulation, predominantly around the leaf margins (Fig. 1A). This is in marked contrast to the pattern previously observed in the parental line (Curtis et al., 2013), where GUS expression is restricted to the midvein and only after exposure to cJ for >16h (Fig 1A). In both cases, the mutation segregated 3:1 with respect to the GUS phenotype suggesting that it was a single recessive mutation. To identify the underlying mutation in line 48_67, we generated a mapping population by crossing a homozygous line to the Arabidopsis ecotype *Ler*. By combining microsatellite mapping of the F2 population with F2 bulk segregant analysis and whole genome resequencing of pools, we identified a region on chromosome 3 as likely being causal for the misexpression phenotype and linked the phenotype to SNPs in 5 putative candidate genes. Further resolution of the mutations with CAPS markers specific for each candidate genes eliminated two candidates. For the remaining three candidate genes, respective T-DNA lines were crossed with the pF-Box:GUS line and the F2 population evaluated for GUS mis-expression. Only in the case of a cross with line Salk_111245, which harbours a T-DNA insertion in gene At3g06290, could we identify F2 individuals with an identical ectopic GUS expression pattern as observed in line 48_67 (Fig 1C). In Salk_111245 the T-DNA insertion localises to the last exon, exon19, of At3g06290. The genomic region of At3g06290 in line 48_67 was amplified by PCR and sequencing confirmed the presence of a point mutation in the seventh exon of the gene, a C to T transition which converts tryptophan residue 263 into a premature stop codon and causes a large deletion in the predicted open reading frame, leading to a loss of over 80% of the predicted polypeptide (Figure 1C). At3g06290 encodes a large gene which comprises 19 exons and encodes a protein of 1697 amino acids which shows similarity to the yeast nuclear pore complex (NPC) protein SAC3. In yeast, SAC3 is part of the TREX-2 (transcription-coupled export-2) complex, a nuclear pore complex implicated in a spectrum of biological roles including preventing genome instability, transcription initiation, gene-gating, and mRNA export (Stewart, 2019). Previous studies have identified that in Arabidopsis two additional genes besides At3g06290 contain the conserved middle region of the yeast SAC3: At2g39240 and At3g54380; At3g06290 has therefore been designated SAC3B, with At2g39240 being named SAC3A and At3g54380 named SAC3C respectively (Lu et al., 2010; Yang et al., 2017).

**Figure 1:**
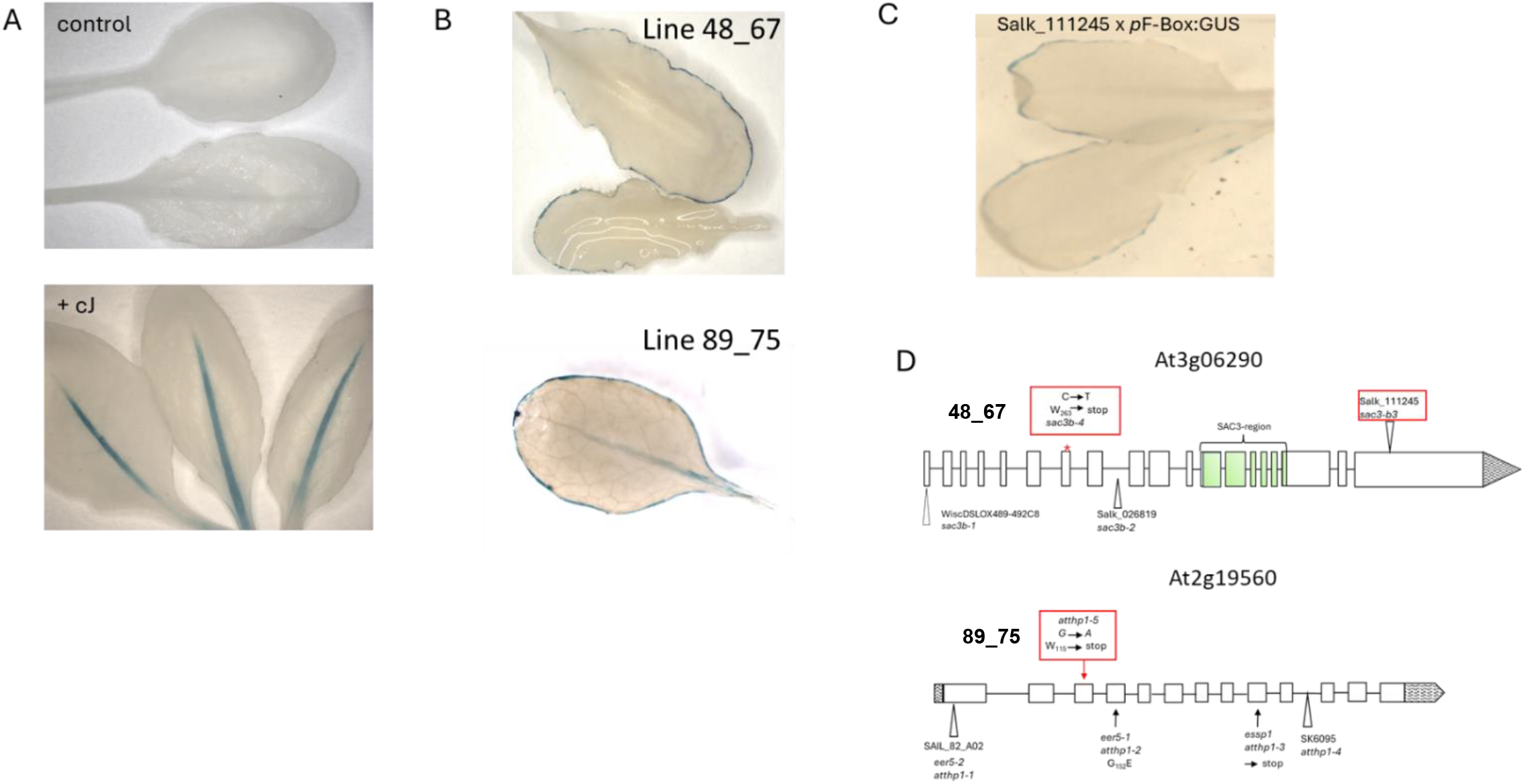
Mutations in components of the Arabidopsis Trex-2 complex are responsible for the deregulation of the expression of the *p*F-Box:GUS reporter. (A) After exposure to the volatile cJ expression of GUS controlled by the KFB39 promoter (Kelch-like domain F-box, At2g44130) is restricted to the leaf mid-rib of mature rosette leaves. (B) In the EMS generated mutant lines 48_67 and 89_75 GUS expression occurs in the absence of cJ, however, expression is not detected in the leaf mid rib but is localised to the leaf margins. (C) The replication of the line 48_67-GUS staining pattern in the cross between Salk_111245 line and the *p*F-Box:GUS reporter line confirms that the EMS generated mutant occurred in the Sac3B gene. (D) Scheme of the genomic organisation of the Sac3B (At3g06290) and THP1 (At2g19560) genes. New EMS alleles of these genes generated in this study are highlighted by a red box.

A similar approach as described for the identification of the underlying mutation in line 48_67 was followed to identify the mutation causing the mis-expression of the GUS reporter in line 89_75. We identified a G to A transition in exon 3 of the gene At2g19560 which converted W_115_ to a stop codon thereby severely truncating the predicted open reading frame of the underlying gene. At2g19560 is predicted to have 13 exons and encodes a polypeptide of 413 residues – thus, the mutation in line 89_75 removes ∼75% of the deduced amino acid sequence (Figure 1D). At2g19560 has previously been identified as a homolog of the yeast gene THP1, which, like SAC3, also is a component of the TREX-2 complex (Lu et al., 2010; Stewart, 2019). In yeast, TREX-2 is a complex comprised of 4 components, Sac3, Thp1, Sus1 and Cdc31, which is tethered to the inner side of the NPC via the nucleoporin Nup1 (Fischer et al., 2002). The Sus1 component interacts with the SAGA transcriptional co-activator complex and the TREX-2 complex has been proposed to functionally couple SAGA-dependent transcription to mRNA export at the inner side of the NPC (Rodríguez-Navarro, 2012). Thus, in our screen for mutants de-regulated in the expression of the KFB39 promoter in the absence of the oxylipin inducer cJ, we identified two different components belonging to the TREX-2 complex, invoking the actions of this nuclear complex in the regulation of the F-Box promoter-reporter used in our study.

### Phenotypes of TREX-2 Arabidopsis mutants

The mutation in At2g19560 in line 89_75 is an additional and new mutant allele of THP1 and, in accordance with the current labelling, was designated *atthp1-5* (Figure 1D). Mature *atthp1-5* plants displayed reduced growth with a more compact shape and smaller, curly leaves. This phenotype was similar to the one observed for previously characterised T-DNA mutant alleles of At2g19560 such as SAIL_82_A02, also designated *atthp1-1* (Lu et al., 2010), and *eer5-1/atthp1-2* (enhanced ethylene response5) (Christians et al., 2008). Interestingly, and contrary as to described by Lu et al., 2010 who did not observe a phenotype in two lines harbouring independent T-DNA KO alleles of SAC3B, (line WiscDSLOX489-492C, allele *atsac3b-1* and line Salk_026819, allele *atsac3b-2;* Figure 1D) T-DNA insertion lines Salk_111245 and Salk_065672 and our EMS generated mutant collectively showed the same morphological differences compared to WT: rosette sizes at 2 weeks were noticeably reduced, leave margins were characteristically folded inwards and the stature of the mature plant was reduced (Figure 2B). This curling of the leaf and the fact that GUS expression in the mutant is restricted to the leaf margin prompted us to investigate whether morphological defects can be linked to this phenotype. Leaf margins of mature WT and Salk_111245 leaves were analysed by Scanning EM (Supplementary Fig S1), however we could not observe a difference in the morphology of the typically elongated leaf margin cells between the two genotypes. However, we did notice that the trichomes of the Sac3B KO line frequently have more solid stalks and are more branched than their WT counterparts (Supplementary Figure S1B). In line with current nomenclature, line Salk_111245, used here to confirm the ectopic expression of the KFB39 promoter, and our EMS generated mutant 48_67 were defined as new alleles *atsac3b-3* and *atsac3b-4* of Sac3B respectively. Data from Lu et al. (2010) showed that AtSAC3B and AtSAC3A are both part of the TREX-2 complex, however, no interaction of SAC3C with other components of this complex could be confirmed. As we have identified new SAC3B mutant alleles which are associated with a morphological phenotype we proceeded to generate *sac3a/sac3b* and *sac3b/sac3c* double mutants and the *sac3a/sac3b/sac3c* triple mutant in order to confirm or reject the hypothesis that different SAC3 members have roles in the same pathway. Arabidopsis T-DNA insertion lines described in Lu et al., 2010 for AtSAC3A (SAIL584_A02) and AtSAC3C (Salk_083805) were obtained and crossed to *atsac3b-3* (Salk_111245). We chose the T-DNA allele of SAC3B instead of the EMS generated mutant as parent in order to avoid introducing any residual additional background mutations associated with our EMS treatment. Whereas previously none of the single and double *atsac3* mutations were reported as displaying a morphological mutant phenotype (Lu et al., 2010), this was not the case in the present study. Specifically, using *atsac3b-3* as parental allele, both double mutants *sac3a/sac3b* (Figure 2C) and *sac3b/sac3c* (Figure 2D) were pronouncedly dwarfed and developed bushy rosettes with small, curly leaves and significantly stunted inflorescences which developed only a few siliques containing viable seeds. These additive effects of mutated SAC3A or SAC3C with *atsac3b-3* suggest that the different SAC3 family members play overlapping roles in the same pathway. The two double mutants were then crossed to generate a triple mutant deficient in all three SAC3 genes. The recovered triple mutant *atsac3a/atsca3b/atsac3c* showed an extreme dwarf phenotype, inflorescences did not develop which precluded setting of seeds (Figure 2E). Therefore, a triple KO line needed to be freshly generated from a parent in which either SAC3A, SAC3B or SAC3C had remained heterozygous.

**Figure 2:**
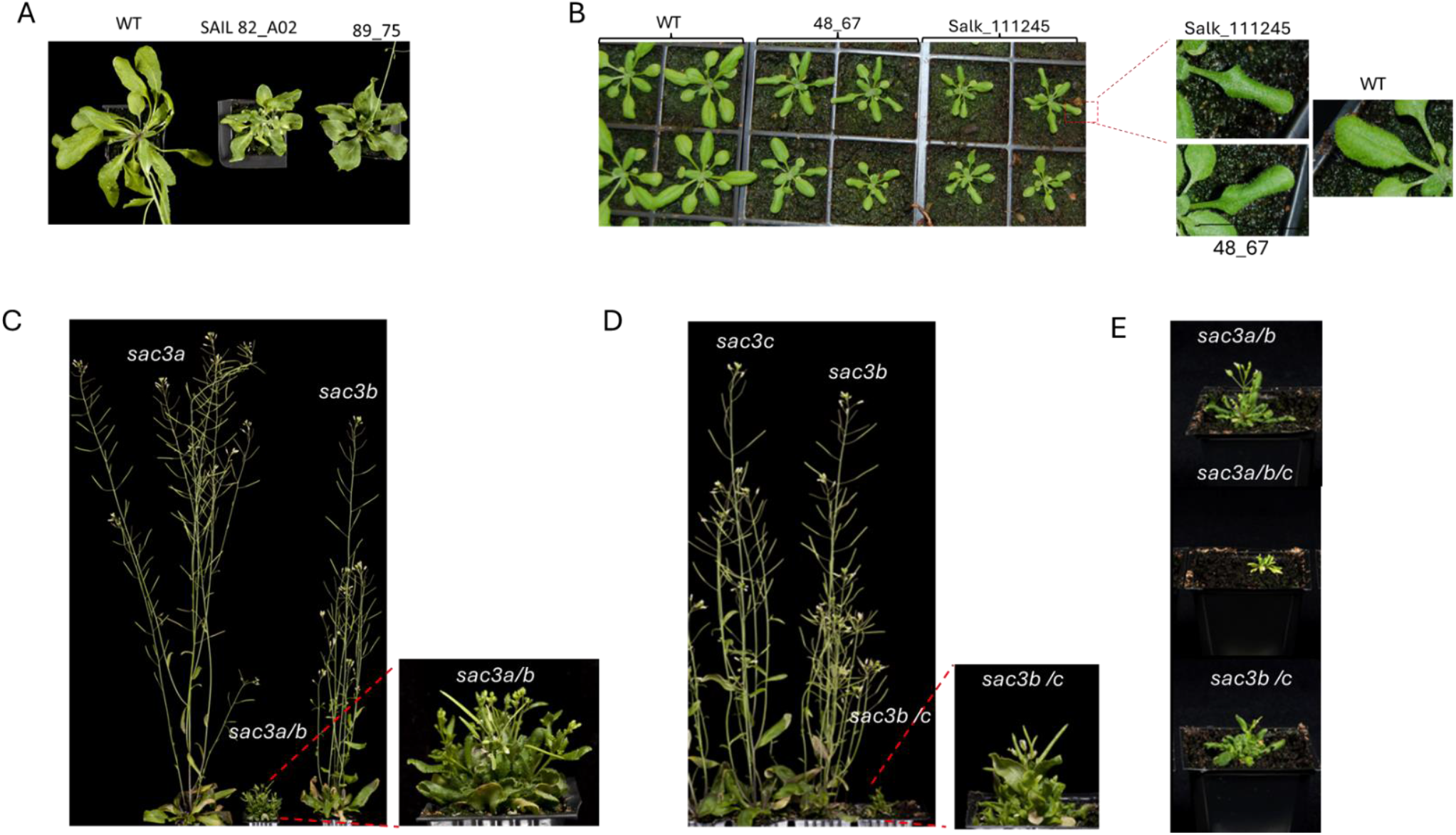
Mutations in components of the plant Trex-2 complex affect plant morphology and their additive effects become evident by progressively more severe phenotypes in double or triple mutants. (A) The EMS generated mutation of the THP1 component of the Trex-2 complex in line 89_75 generates plants with reduced growth and a more compact rosette with smaller, curly leaves. This is similar to the phenotype observed for the THP1 knockout line SAIL82_A02 harbouring the *atthp1-1* allele for this gene (Lu et al., 2010). (B) The EMS generated mutation of Sac3B in line 48_67 and the respective knockout allele in line Salk_111245 cause a reduction in rosette sizes and a characteristic folding of the leaf margins. (C) A double homozygous knock out line for Sac3A and Sac3B components of the Trex-2 complex is a bushy dwarf with reduced leaf laminae. Inflorescences do not elongate but carry flowers which are capable of developing siliques containing viable seeds. A similar phenotype is observed when both, Sac3C and Sac3B, are disrupted (D). Images were taken of mature 5 weeks old plants grown in long day conditions. The most severe phenotype is shown by a *sac3a/sac3b/sac3c* triple knock out line which has lost the ability to develop siliques and needs to be generated *de novo* from a parent in which either Sac3A, Sac3B or Sac3C had remained heterozygous (E).

Taken together, the additive nature of these mutations suggests that all three SAC3 proteins are very likely involved in the same pathway which, analogous to the role of SAC3 in yeast, is presumably facilitating mRNA export from the Arabidopsis nucleus through the nuclear pore into the cytoplasm.

### Functional evidence that AtTHP1 and AtSAC3B are involved in mRNA export from the nucleus

To test this hypothesis, we determined whether mRNA export was compromised in plants harbouring mutations in the distinct components of the putative Arabidopsis TREX-2 complex which we have identified in our screen, by examining the localization of poly(A) mRNA in young leaves using *in situ* hybridization (Gong et al., 2005).

In accordance with data reported for the mutant in the AtTHP1 component of the TREX-2 complex, *atthp1-3*, (Lu et al., 2010) we could show that, compared with wild-type, poly(A) mRNA preferentially accumulated in the nuclei of our EMS mutagenized line 89_75, the *atthp1-5* mutant allele, suggesting that mRNA export is compromised (Figure 3A) in this line. The finding that AtTHP1 is necessary for nuclear mRNA export supports the hypothesis that it is a component of the putative Arabidopsis TREX-2 complex.

**Figure 3:**
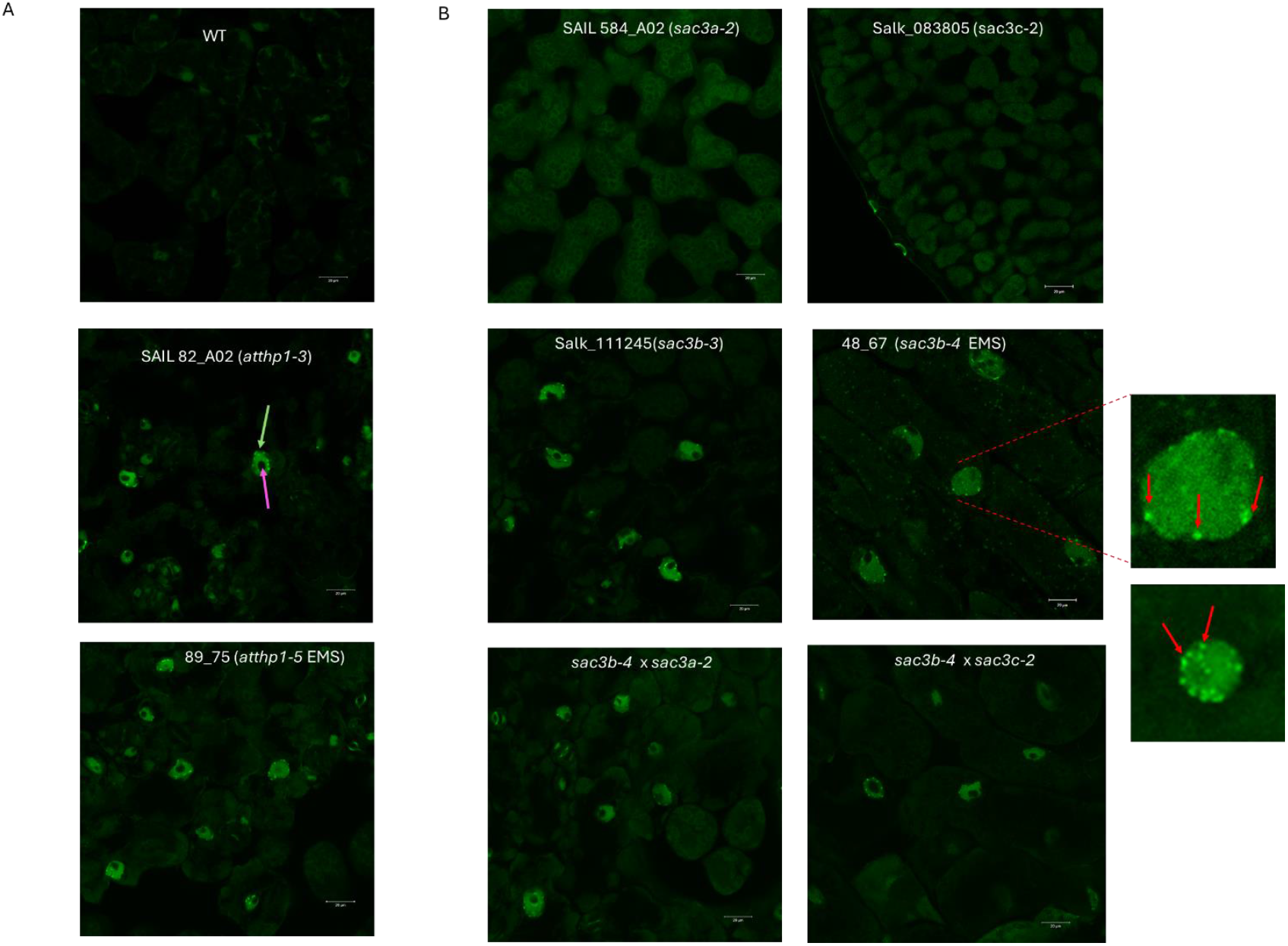
mRNA export is inhibited when the THP1 or Sac3B components of the Trex-2 complex are non-functional. (A) polyA mRNA accumulates in the nucleus when THP1 function is disrupted either by T DNA insertion (SAIL82_A02) or by point mutation as seen in line 89_75 isolated from our EMS screen. Yellow arrow: nucleus; magenta arrow: nucleolus. Bar = 20 μm. (B) The Sac3B component of the Trex-2 complex is crucial for polyA mRNA export from the nucleus to proceed normally whereas Sac3A or Sac3C components play less significant roles. In *sac3a* and *sac3c* knock out lines polyA mRNA is retained in the nucleus only when *sac3b* is defective as observed in single and double knockout lines respectively. Nuclei with inhibited mRNA export show foci of high GFP intensity, possibly forming at the nuclear envelope (red arrows), where polyA mRNA accumulates, however the significance of which remains unknown. Bar = 20 μm.

Concerning the SAC3 components of the TREX-2 complex, Lu et al. (2010) did not detect any retention of poly(A) mRNAs either in the nuclei of SAC3A, SAC3B or SAC3C single mutants or their respective double or triple mutants. Our poly(A) mRNA hybridisation assay performed on single SAC3A *(sac3a-2* allele) and SAC3C *(sac3c-2* allele) mutants confirmed their observation that in these mutants, mRNA export from the nucleus proceeds normally. However, unlike what was observed by Lu et al. (2010), poly(A) mRNA accumulated in the nuclei of both our new alleles for *sac3b*, irrespective of whether it was EMS generated *(sac3b-4* mutant line 48_67) or a T-DNA disruption allele *(sac3b-3)* suggesting that the SAC3B component of the TREX-2 complex is required for mRNA to be efficiently exported from the nucleus (Figure 3B). In agreement with this, *sac3* double mutants in which the SAC3B component is defective also retained mRNA in their nuclei. Interestingly, we noticed that nuclei of mutants with a non-functional SAC3B component showed bright fluorescent loci at the nuclear envelope which indicates high concentration of polyA mRNA at these positions (Figure 3B), however at this stage the significance of this observation remains unknown.

Taken together, our results suggest that AtTHP1 and AtSAC3B actively facilitate the export of mRNA from the nucleus through the nuclear pore into the cytoplasm whereas SAC3A and SAC3C are not directly involved. Put into context with our observation that SAC3A/SAC3B and SAC3B/SAC3C double mutants have a significantly more severe morphological phenotype than the SAC3B single mutant raises the interesting question if and how SAC3A and SAC3C are contributing to efficient mRNA export.

### Defects in Thp1 and SAC3B increase ethylene sensitivity in seedlings

The <u>*e*</u>nhanced <u>*e*</u>thylene <u>*r*</u>esponse *eer5-1* mutant *(atthp1-2* in Lu et al., 2010) has been isolated in an EMS screen for extreme hypocotyl shortening in the presence of saturating ethylene concentrations and is due to a G152E substitution in exon 4 of the Thp1 component of the Arabidopsis TREX-2 complex (Christians et al., 2010). The increased sensitivity to ethylene in *eer5* was suggested to be associated with the loss of expression of a subset of genes rather than increased expression (Christians et al., 2010; Deslauriers et al., 2015). We therefore analysed whether this enhanced response to ethylene is uniquely associated with a defective Thp1 subunit of TREX-2 or whether SAC3 components also play a role in this response, by germinating seedlings on plates containing the immediate ethylene precursor 1-aminocyclopropane-1-carboxylic acid (ACC) and measuring hypocotyl length after 4 days of growth in the dark. We found no difference in ACC-induced inhibition of hypocotyl elongation in WT, *atsac3a-2* and *atsac3c-2* single mutants suggesting that SAC3A and SAC3C are not involved in mediating this response (Figure 4A). Confirming previous results (Christians et al., 2010), seedlings of line 89_75 harbouring our new Thp1 mutant allele *atthp1-5* exhibited a significantly enhanced response to ACC (Figure 4B). We also detected extreme hypocotyl shortening in the presence of ACC in seedlings with mutant alleles of SAC3B, line 48_67 *(sac3b-4)* and Salk_111245C *(sac3b-3)* which strongly suggests that SAC3B is also involved in mediating this response.

**Figure 4:**
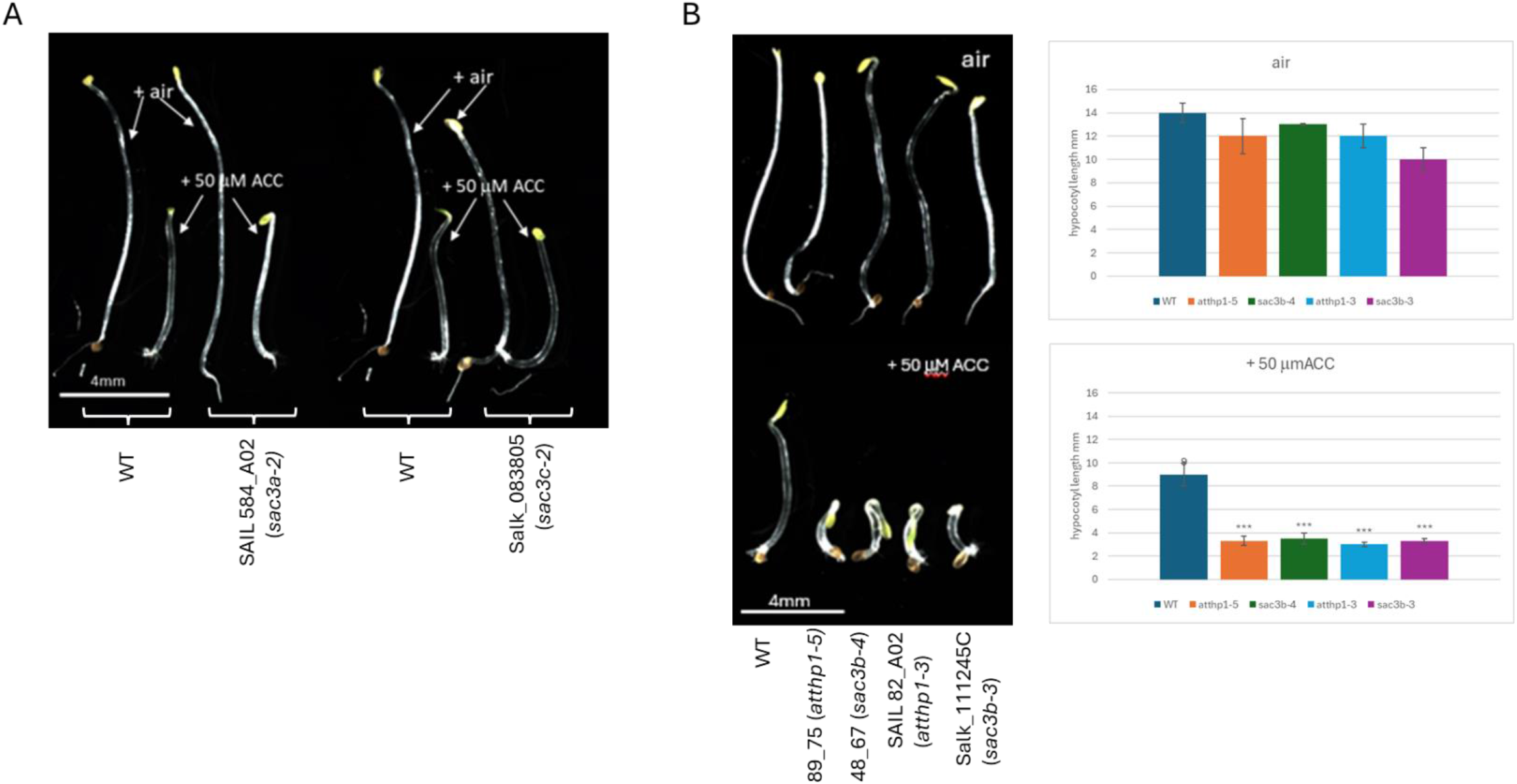
THP1 and Sac3B but not Sac3A and Sac3C single mutants exhibit enhanced sensitivity to ethylene. (A) Ethylene response phenotype of *sac3a* and *sac3c* single mutants is similar to the ethylene response observed in WT whereas the single mutants of *atthp1* and *sac3b* exhibit enhanced sensitivity to ethylene (B). Scale bar: 4mm.

### Expression of AtSAC3 genes

The *p*F-Box :GUS reporter showed a very discrete pattern of expression not only when misexpressed as a consequence of mutations in components of the TREX-complex but also in the presence of the xenobiotic inducer (Curtis et al., 2013). Therefore, we wished to determine whether expression patterns of TREX-2 components could be implicated in the regulation of the F-Box gene. Promoter-GUS fusions were generated with promoters from AtSAC3A, B and C respectively and GUS expression evaluated for independent transgenic events in Arabidopsis. When driven by the AtSAC3B promoter, GUS staining was detected in the vasculature of the leaf, partially overlapping with the observed F-Box KFB39 expression; however, this extended further into the veins beyond the midrib (Figure 5e). In addition, GUS staining was observed in pollen grains and in the abscission zone of floral organs, both of which were not detected in the cJ-induced KFB39-GUS line (Figure 5f, g, h). Using the AtSAC3A promoter, GUS staining was detected in the leaf and sepal vasculature but seemed to be absent in the petal vascular system (Figure 5a and b). Like AtSAC3B, AtSAC3A is expressed in pollen grains (Figure 5c). Interestingly, we found that SAC3A expression is highly inducible by wounding as leaves detached with scissors or damaged by forceps showed intense staining around the injured tissue (Figure 5d). AtSAC3C expression in mature leaves was predominantly present in the hydathodes (Figure 5i). In floral organs we detected strong GUS staining in the style, ovules and pollen grains (Figure 5j and k). Close up of the ovules revealed that GUS staining was restricted to the embryo sac (Figure 5l). Collectively, these data indicate very limited overlap in the native expression of F-Box KFB39 and the AtSAC3 genes.

**Figure 5:**
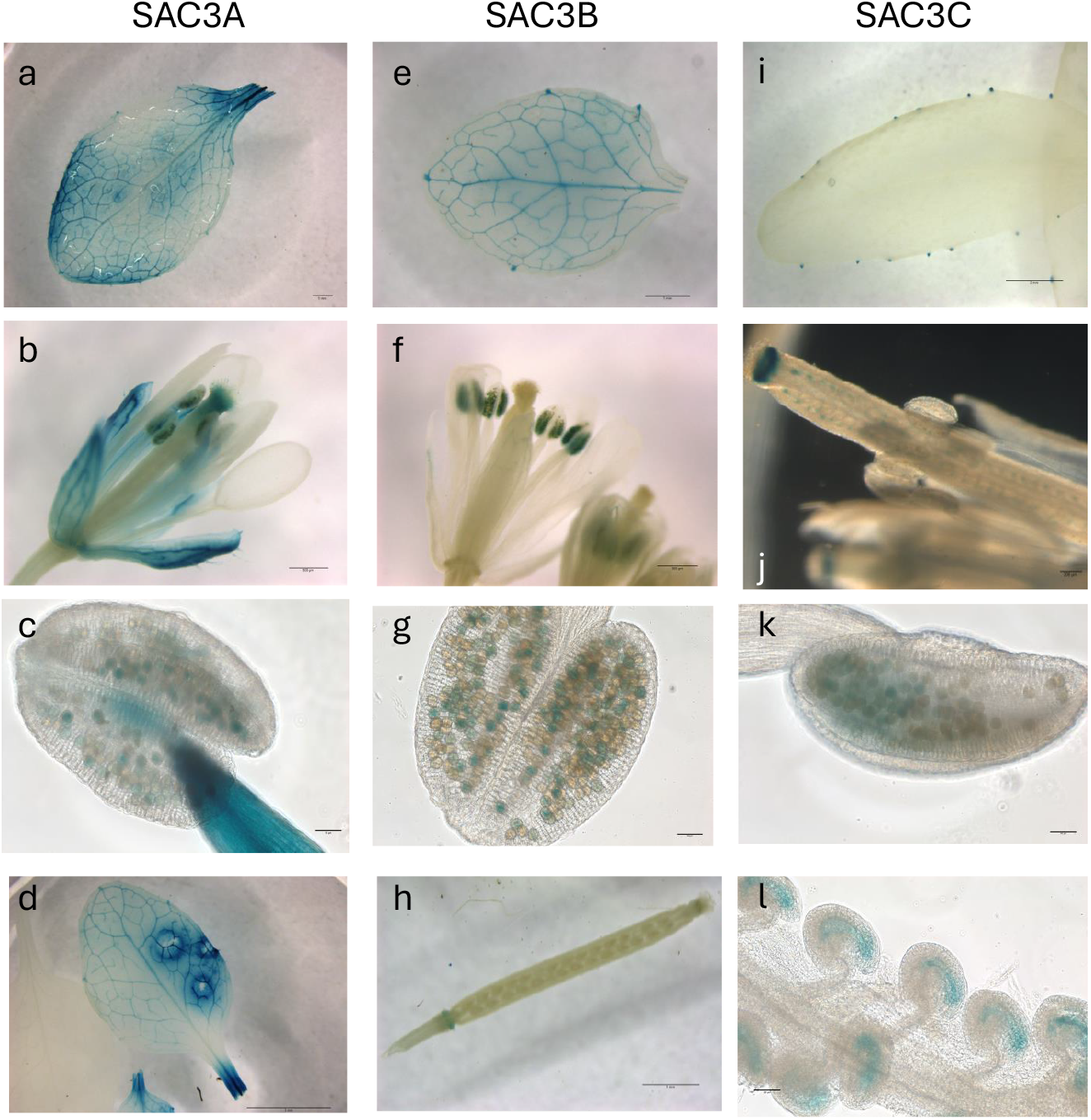
Promoter:GUS reporter assays for determination of tissue specific expression of Sac3A, Sac3B and Sac3C respectively. A 1.27 kb promoter fragment for Sac3A was used to drive GUS expression. GUS staining was observed in mature leaves with intense staining concentrated around the vasculature (a; bar = 1mm) whereas in flowers staining was observed in the sepal vasculature, style, anthers and pollen (b, c; bar = 500μm and 50μpm respectively). Sac3A expression is sensitive to wounding as we noticed that areas surrounding injured tissue showed high intensity staining (d; bar = 3mm). Sac3B expression was observed in mature rosette leaves mainly in vascular tissue (e; bar = 1mm) and in pollen (f, g; bar = 500μm and 50μm respectively). GUS staining was also observed in the dehiscence area of floral organs around the base of siliques (h; bar = 1mm). Mature rosette leaves of a line harbouring a 0.92 kb Sac3C promoter:GUS reporter fusion showed GUS staining localised to hydathodes (I; bar = 2mm). In floral organs strong staining was detected in the style, some pollen grains and in ovules (j, k; bar = 200μm and 50μm respectively). Higher magnification of the ovules could localise GUS expression to the embryo sac of the ovules (I; bar = 50μm).

## Discussion

The isolation of mutant alleles of AtSac3B and AtTHP1 has previously been reported on a number of occasions, from apparently diverse and unrelated screens, making it intuitively challenging to define the contribution these genes might play in a genetic pathway for the regulation of KFB39. Previously, these two components of the TREX-2 complex have been hypothesised to be involved in the regulation of transgenes and epigenetic control, perhaps through the export of miRNAs. The first report of the isolation of a mutation in TREX-2 in Arabidopsis (Lu et al., 2010) was based on a screen for mutants that allowed ectopic expression of the 2S seed storage protein (Tang et al., 2008) – the recovered mutant *(essp1:* ectopic expression of seed storage protein) was subsequently shown to have a stop mutation in the 9^th^ exon of AtTHP1 and hence renamed *atthp1-3*. Intriguingly, the mis-expression of the 2S seed storage protein promoter-GUS fusion in the *atthp1-3* mutant is apparently very similar (Supplementary Table S1) to that we observed in this study with the KFB39-GUS reporter in either the *atsac3b* or *atthp1-5* mutant described here. However, there is no obvious commonality between these two reporters constructs other than the fact they are both using GUS and contain a nopaline synthase 3’ UTR and terminator. There are no shared elements in the 2S seed storage protein and KFB39 promoters and they display very different native expression patterns, with the 2S seed storage protein being exclusively seed-specific whereas the KFB39 is expressed in photosynthetic tissue but only in the presence of the volatile oxylipin *cis*-jasmone. Thus, it is not obvious why disruption of components of the TREX-2 complex would lead to a shared deregulated, mis-expression pattern, unless this complex was responsible for the synthesis of factors (such as miRNAs or repressors) that impact on the regulation of these promoters.

Intriguingly, the identification of mutations in TREX-2 components have arisen multiple times from forward genetic screens based on the deregulated expression of a transgene promoter-reporter construct. These current known examples, summarised in Supplementary Table S1, include 2S seed storage protein *essp1* (described above), as well as our deregulated expression of KFB39. Other examples include loss of promoter activity, such as the activity of the RAD51 promoter. *RAD51* expression is induced by genome instability and endoreplication, and GFP expression from this promoter is elevated in the *atxr5/6* background. Suppressors of *atxr5/6* were identified by reduced expression of GFP in their cotyledons and two mutants were in the TREX-2 complex in AtSAC3B and AtTHP1 (Hale et al., 2016). A similar forward genetic screen was used to identify anti-silencing factors which might modulate the expression of viral-expressed reporters (double CaMV35S-LUX and double CaMV35S-NPTII) (Yang et al., 2017). These transgenes were expressed in the *rdr6-11* mutant background, and a line which showed reduced luciferase expression was found to be mutated in AtSAC3B. There is no obvious relationship between *atxr5/6* and *rdr6-11*, nor between the promoter-reporter constructs used in these studies, but both resulted in the isolation of mutant alleles of *atsac3b*. More recently, an EMS forward genetic screen to identify components involved in miRNA biogenesis and/or activity, was performed on the pSUC2: amiR-SUL (amS) line25. amS expresses the amiR-SUL artificial miRNA from the SUC2 promoter, which is specific for phloem companion cells (Zhang et al., 2020). Silencing of SULFUR (SUL, also known as CHLORINA42, a gene required for chlorophyll synthesis26) by amiR-SUL causes bleaching along the leaf veins. A mutant with reduced leaf bleaching, a phenotype indicative of compromised amiR-SUL activity, was shown to be a new allele of *atthp1*, implicating TREX-2 in this process (Zhang et al., 2020). Using some of the earlier identified mutant alleles of the three AtSac3 genes and another component of TREX-2 (NUP1), a role in the accumulation of miRNAs for this complex was proposed (Zhang et al., 2020).

Collectively, these mutant screens, although very interesting, complicate our understanding of the role of the TREX-2 complex in Arabidopsis since they invoke it in a range of apparently unrelated processes (miRNA accumulation, anti-silencing, genome stability, tissue-specific expression). The general model for how TREX-2 functions is that it serves to connect the SAGA and MEDIATOR complexes to the nuclear pore, possibly delivering “gene gating” (which might also include miRNA and other non-coding sequences). In the specific case of the deregulation of the *cis*jasmone-induced KFB39 promoter, the involvement of TREX-2 is perplexing but there are some possible explanations for our observations. Until relatively recently, the substrates for Kelch-domain F-Box proteins such as KFB39 were unknown, but it is now clear that these proteins are involved in the targeted proteolysis of the enzyme phenylalanine ammonia lyase (PAL) (Zhang et al., 2013), the first and committed step in the phenylpropanoid pathway and is therefore involved in the biosynthesis of the polyphenol compounds such as flavonoids, phenylpropanoids, and lignin in plants (Kim et al., 2020). Several KFBs, including KFB39, were shown to specifically modulate the accumulation of PAL (Zhang et al., 2015), although a direct role for this enzyme in *cis*-jasmone mediated responses has not previously been observed. Interestingly, we previously reported that modulating the expression of KFB39 altered the colonization of Arabidopsis by root knot nematodes (Curtis et at 2013), and we now believe that this occurred as a consequence of varying the levels of flavonoid-derived components that presumably act as cues or stimuli to the RKNs. Recently, Wang et al (2022) carried out comprehensive transcriptome analyses of Arabidopsis mutants involved in phenylpropanoid metabolism, confirming a central role for KFB39 in that pathway. Intriguingly, expression of KFB39 is strongly reduced in *med5a/b* mutants in the MEDIATOR complex – as noted above, the MEDIATOR complex interacts with TREX-2 and SAGA PolII (Dolan and Chapple, 2017), providing some additional independent links between TREX-2 and the regulation of the KFB39 promoter. Similarly, the transcriptome of MED5-modulated pathways (Kim et al., 2020) shows some overlap with the transcriptome we previously reported for *cis*-jasmone-induced gene expression (Matthes et al., 2010).

### Other possible scenarios to explain the involvement of TREX-2 in diverse processes in Arabidopsis

A frustrating consequence of considering all of the studies listed above which have resulted in the isolation of Arabidopsis mutants defective in TREX-2 components is the lack of any obvious commonality between all the different studies (listed in Supp Table S1). Equally, it is striking that mutants in TREX-2 are repeatedly recovered from forward genetic screens based on the deregulated expression of promoter-reporter constructs – an explanation for this is currently lacking. However, it is possible to propose some scenarios which might link the various studies and as such, these could form the basis for further research. Possible explanations include:

- The loci for T-DNAs carrying the promoter-reporter constructs are all perturbing a common pathway or node in the regulation of the TREX-2 complex. Such a scenario seems unlikely, but in the absence of data remains a possibility. Currently, there is no information as to the genomic locations of any of the promoter-reporter constructs used to recover mutant alleles of TREX-2.
- That miRNAs are involved in the regulation of all the promoters under study and their biogenesis/export is disrupted in the *trex-2* mutants. This also seems unlikely since some of the promoters used in these studies are very well characterised.
- There are common elements/sequences present in all the promoter-reporter constructs – for example, several cassettes contain the nopaline synthase 3’ UTR. The presence of the NOS 3’-UTR represents the strongest commonality amongst all the promoter-reporter constructs. However, it is not present in all of them e.g. Zhang et al., 2020.
- Some other epigenetic process that was invoked by the delivery of the promoter-reporter transgenes results in reporter misexpression/deregulation.
- Mutations in TREX-2 result in the nuclear retention of common repressors or related TFs that modulate the transcription of the promoters used in these different studies. However, one might expect that if this was the case, then mutant alleles of TREX-2 components might be recovered from other forward genetic screens.

## Conclusions

*We* have recovered two new alleles *(atsac3b-4, atthp1-5)* of components in the TREX-2 complex that result in the mis-expression of the *cis*-jasmone-responsive KFB39 promoter. KFP39 is a Kelch-domain F-box protein that is responsible for the proteolysis of PAL, the first step in the biosynthesis of phenylpropanoids. This study implicates the TREX-2 complex, a presumptive component of the nuclear pore complex, in this process. Mutant alleles of SAC3 and THP1 have been recovered from multiple unrelated screens, with the only common factor being selection based on mis-expression of a promoter-reporter transgene. Based on these collective observations, we conclude that in Arabidopsis TREX-2 plays a currently unresolved role in gene expression.

## Acknowledgements

This study was partially supported by BBSRC grants BBS/E/C/00005207 and BBS/E/C/000I0420.

## Supplementary Data

**Supplementary Figure S1** – Scanning EM of *atsac3b*

**Supplementary Table S1** – Summary of different promoter-reporter screens which have recovered mutant alleles in TREX-2

