## Supplementary material for "Mutations in components of the TREX-2 complex result in misexpression of the Kelch-domain F-Box protein KFB39 promoter in *Arabidopsis thaliana*": Fig S1

### Slide 1
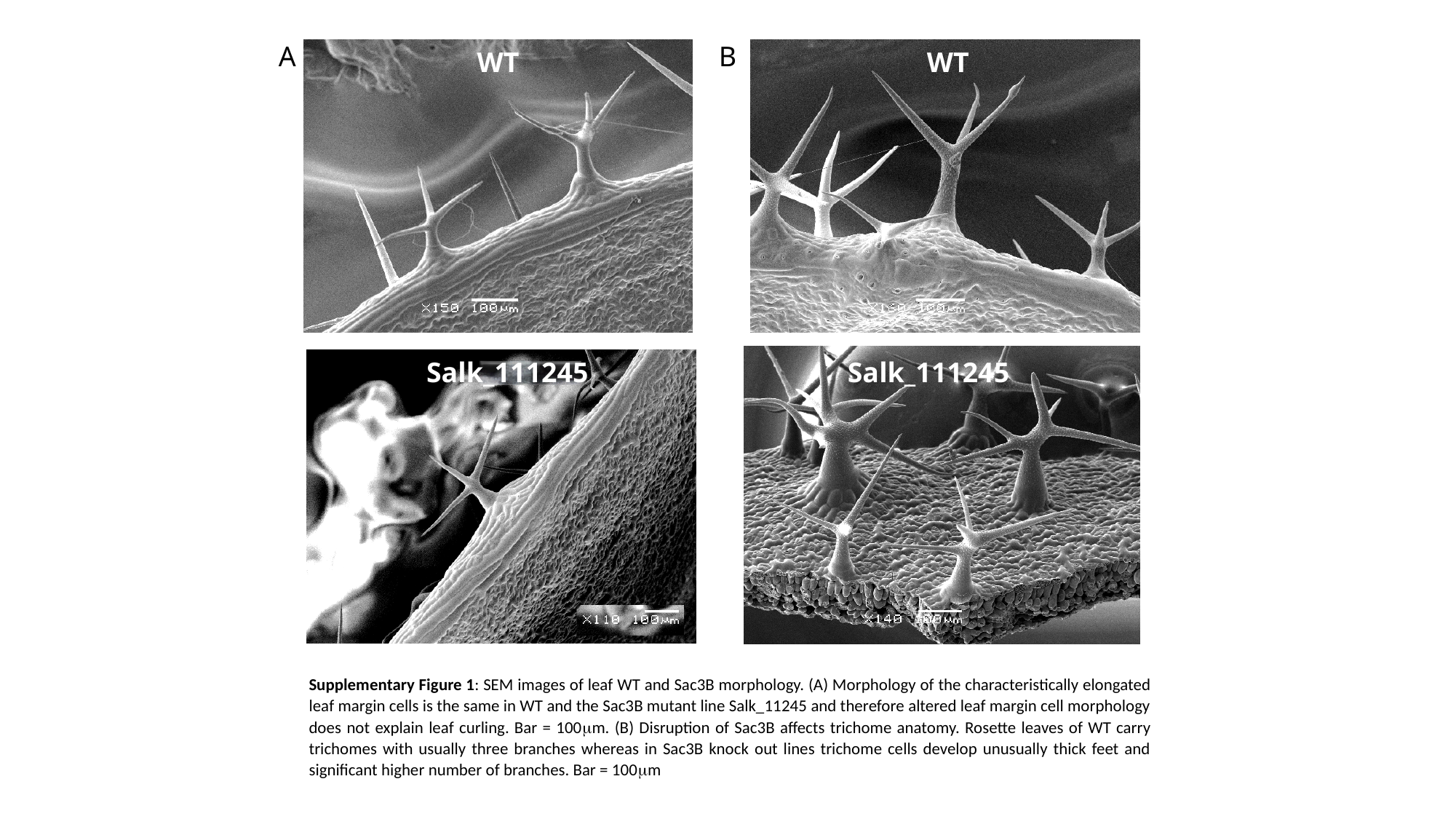

A
B
WT
WT
Salk_111245
Salk_111245
Supplementary Figure 1: SEM images of leaf WT and Sac3B morphology. (A) Morphology of the characteristically elongated leaf margin cells is the same in WT and the Sac3B mutant line Salk_11245 and therefore altered leaf margin cell morphology does not explain leaf curling. Bar = 100μm. (B) Disruption of Sac3B affects trichome anatomy. Rosette leaves of WT carry trichomes with usually three branches whereas in Sac3B knock out lines trichome cells develop unusually thick feet and significant higher number of branches. Bar = 100μm
