## Supplementary material for "Mutations in components of the TREX-2 complex result in misexpression of the Kelch-domain F-Box protein KFB39 promoter in *Arabidopsis thaliana*": Supp Table S1

**Supplementary Table S1**

| **Gene identified** | **Reporter construct** | **Mutant name** |  | **Reference** |
| --- | --- | --- | --- | --- |
| Thp1, Sac3B | KFB39 (At2g44130) promoter -GUS reporter | 89_75  48_67 | Expression of KFB39pro::GUS in the absence of the inducer cis-jasmone  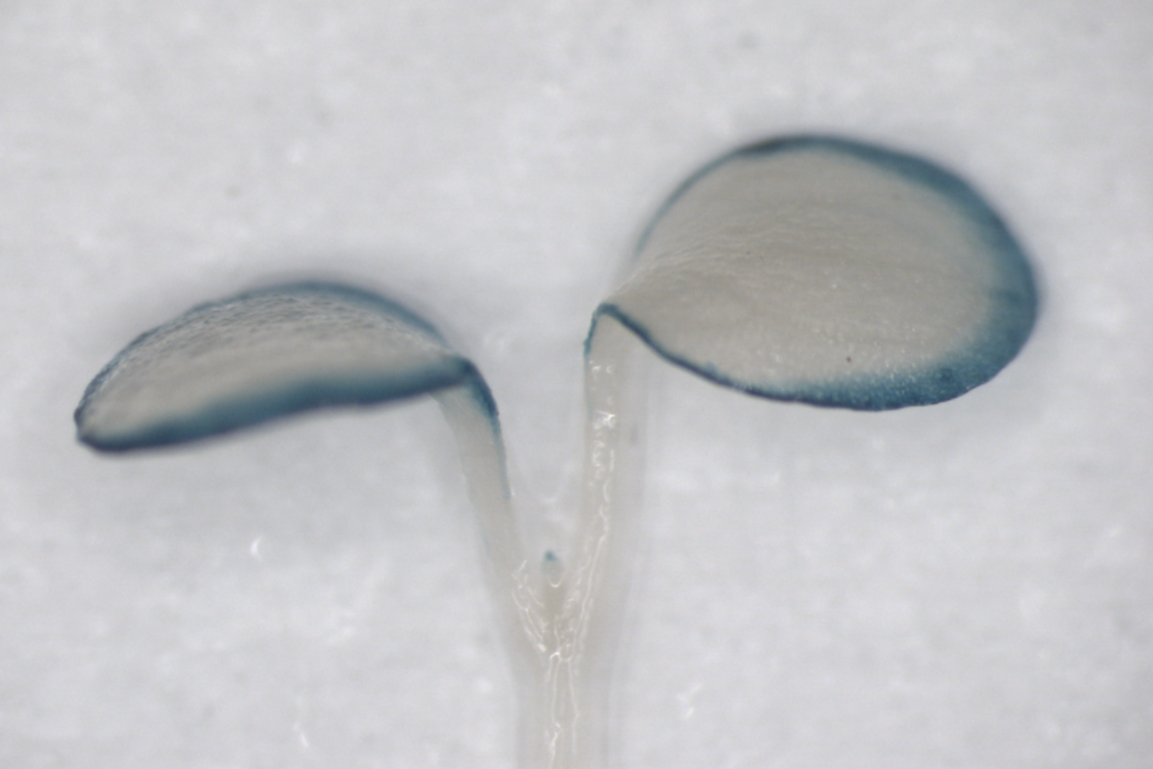 | This study |
| Thp1/ At2g19560 | Ectopic expression of a soybean conglycinin (7S storage protein) gene promoter–GUS transgene (*βCG_pro_:GUS*)  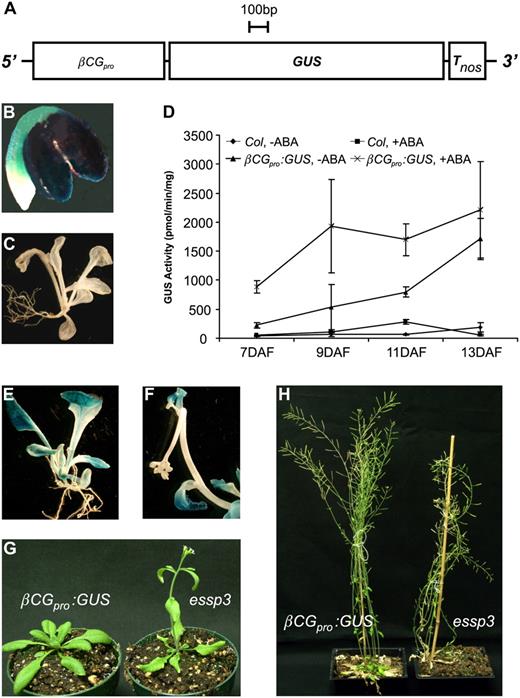 | Essp1 | 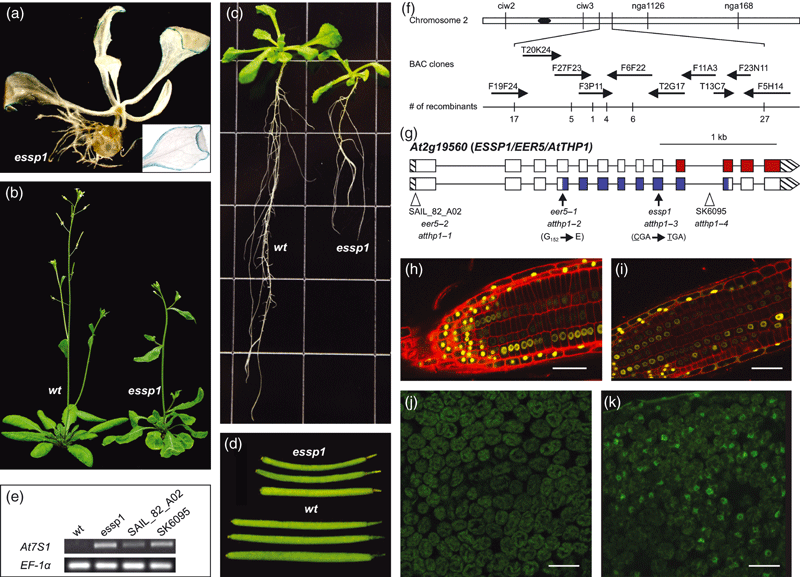 | [**https://doi.org/10.1111/j.1365-313X.2009.04048.x**](https://doi.org/10.1111/j.1365-313X.2009.04048.x)  Lu et al TPJ 2010 |
| Thp1,  Sac3B (At3g06290) | atxr5/6 RAD51pro::GFP | ems_2_37 ems_2_300 | Rad51 expression is induced by genome instability including increased transposon activity and endoreduplication. Mutations in TRex-2 switch off that response, as measured by GFP in the *atxr* background | <https://doi.org/10.1371/journal.pgen.1006092>  Hale et al PLOS Genetics 2016 |
| Sac3B | Transgenic plant previously named YJ, which contains two transgenes, d35S::LUC and d35S::NPTII, in the rdr6-11 background (DOI: [10.1093/nar/gkv958](https://doi.org/10.1093/nar/gkv958)) | P31 | Mutations in TREX-2 components (sac3b, thp1, nup) cause silencing of transgene reporter | <https://doi.org/10.1093/nar/gkw850>  Yang et al NAR 2016, |
| Thp1 | (EMS) mutagenesis screen using the *pSUC2: amiR-SUL* (*amS*) line[25](https://www.nature.com/articles/s41477-020-0726-z#ref-CR25) | *thp1-5 amS* | amS expresses the amiR-SUL artificial miRNA from the SUC2 promoter, which is specific for phloem companion cells. Silencing of SULFUR (SUL, also known as CHLORINA42, a gene required for chlorophyll synthesis by amiR-SUL causes bleaching along the leaf veins. One mutant with reduced leaf bleaching, a phenotype indicative of compromised amiR-SUL activity was in the gene THP1 | <https://doi.org/10.1038/s41477-020-0726-z>  Zhang et al, Nat Plants 2020 |
